# FerroEnrich: An Interactive web tool for computing Ferroptosis index and gene enrichment

**DOI:** 10.1101/2024.09.25.615075

**Authors:** Munichandra Babu Tirumalasetty, Mayank Choubey, Md Wahiduzzaman, Rashu Barua, Mohammad Sarif Mohiuddin, Qing Robert Miao

## Abstract

Ferroptosis is an iron-dependent form of controlled cell death and is characterized by the formation of lipid peroxides. Understanding gene expressions associated with ferroptosis is critical for determining its function in illnesses and potential therapeutic approaches. Despite its significance, no computational model is currently available to accurately quantify the ferroptosis incident. FerroEnrich is a sophisticated web-based tool built using R Shiny application to enumerate the occurrence of ferroptosis based on the relevant gene expressions. This tool available at https://ferroenrich.shinyapps.io/ferroenrich/ processes the input gene expression file to identify genes that are resistant or prone to ferroptosis, calculates ferroptosis index value with dynamic colored heatmap and gene network plot. This manuscript describes the design, operation and usability of FerroEnrich, including examples and a discussion of its potential impact on ferroptosis research. FerroEnrich is a vital tool for researchers, allowing them to explore and analyze complicated gene expression data related to ferroptosis.

## 1. Introduction

Ferroptosis is a unique type of controlled cell death characterized by iron accumulation and lipid peroxidation and differs from apoptosis, necrosis, and autophagy, and plays a critical in the pathophysiology and development of many diseases including liver diseases, cancer and neurological disorders etc., (Dixon et al., 2012; Stockwell et al., 2017). Ferroptosis is driven by the accretion of lipid peroxides to deadly amounts when cellular antioxidants systems, notably those involving glutathione and glutathione peroxide 4 (GPX4), AIFM2 and GCH1 (Friedmann Angeli et al., 2014). This process is fundamentally distinct from other types of cell death because it involves a metabolic catastrophe caused by iron metabolism and reactive oxygen species (ROS). (Yang et al., 2016). In the context of liver diseases, ferroptosis is double-edged sword. It has therapeutic potential by targeting liver cancer cells that are resistant to chemotherapeutical treatments. Nevertheless, uncontrolled ferroptosis can worsen the pathogenesis of other liver diseases such as cirrhosis by causing excessive tissue damage and inflammation, Balancing the activation and inhibition of ferroptosis in specific types of liver cells is therefore critical for achieving treatment efforts in chronic liver disease management (Jiang et al., 2021; Wu et al., 2021).

Gene expression and regulation of ferroptosis is a complex process is and governed by genes such as TFRC, iron absorption, and GPX4 which are required to detoxify lipid peroxides using glutathione (Chen et al., 2023). Another gene, ACSL4 is a predominant regulator for incorporating poly unsaturated fatty acids and making it more vulnerable to oxidative damage (Doll et al., 2017). Furthermore, the cystine/glutamate antiporter system, particularly through the action of SLC7A11, is essential for maintaining glutathione level, which directly affects the cellular antioxidant capability and susceptibility to ferroptosis (Koppula et al., 2021; Yan et al., 2023; Liu et al., 2022). Transcription factors like p53 and NRF2 affect the expression of these genes, regulating ferroptosis, which can be advantageous or deleterious depending on the disease status (Liu et al., 2020; Anandhan et al., 2020). As an example, in hepatocellular carcinoma, increasing ferroptosis could aid in the elimination of resistant cancer cells, whilst, decreasing ferroptosis in chronic diseases such as cirrhosis may mitigate liver damage and fibrosis (Jiang et al., 2024; Ajoolabady et al., 2023). Understanding and controlling these integrative pathways may lead to tailored medicines that promote or inhibit ferroptosis, opening up new possibilities for treating a variety of liver illnesses (Chen et al., 2022).

Due to the complex gene regulation mechanisms observed in various disease states, there is an urgent need for a computational model that can predict and quantify ferroptosis levels under specific disease conditions based on relevant gene expression data (Jin et al., 2021; Feng et al., 2023). The intricate regulatory networks of ferroptosis, involving multiple pathways, make computational web-tools essential for both prediction and intervention (Sun et al., 2023). To understand the complex relationships that control ferroptosis, web-based tools make it possible to identify the predominant regulators from the intricate and large amounts of gene expression data. The recent advancement in computational web resource, FerrDb is specially designed to improve our knowledge of ferroptosis gene regulation. It is a comprehensive database of both promoting and resisting genes of ferroptosis in various disease conditions with a ferroscore for each gene (Zhou et al., 2020; Zhou et al., 2023). Another web resource, FERRREG, catalogs targets, regulators and drug molecules related to ferroptosis-associated diseases (Zhou et al., 2024).

However, there was no web-tool existed that allowed researchers to directly monitor and assess the ferroptosis levels in a variety of disease situations. This notable gap can be satisfied by our FerroEnrich, the first web-based tool specifically designed to enable researchers to actively analyze and predict ferroptosis susceptibility through an interactive web-based platform. Unlike static databases such as FerrDb, which just store gene information about ferroptosis, FerroEnrich allows users to upload gene expression data and quickly produce a Ferroptosis index value. This score gives a quantifiable measure of the propensity for ferroptosis under diverse settings, providing essential insights that can drive research into disease pathways in which ferroptosis plays an important role. This ability distinguishes FerroEnrich as a novel tool, allowing researchers to undertake real-time and facilitate the development of targeted therapeutic approaches in ferroptosis-related disorders.

## 2. Methods

### 2.1. Tool Development

FerroEnrich is built on the R Shiny framework (Jia et al., 2022), which provides a dynamic, web-based interface for interactive data analysis. This flate forms combines the usage of R script (ver. 4.3.1) and is deployed based on shinyapp.io and dynamic capacity Shiny, allowing users to conduct analysis in user-friendly setting. This integration enables real-time data processing and visualization, resulting in a more intuitive understanding of ferroptosis mechanisms based on gene expression patterns (**Figure 1**).

**Figure 1:**
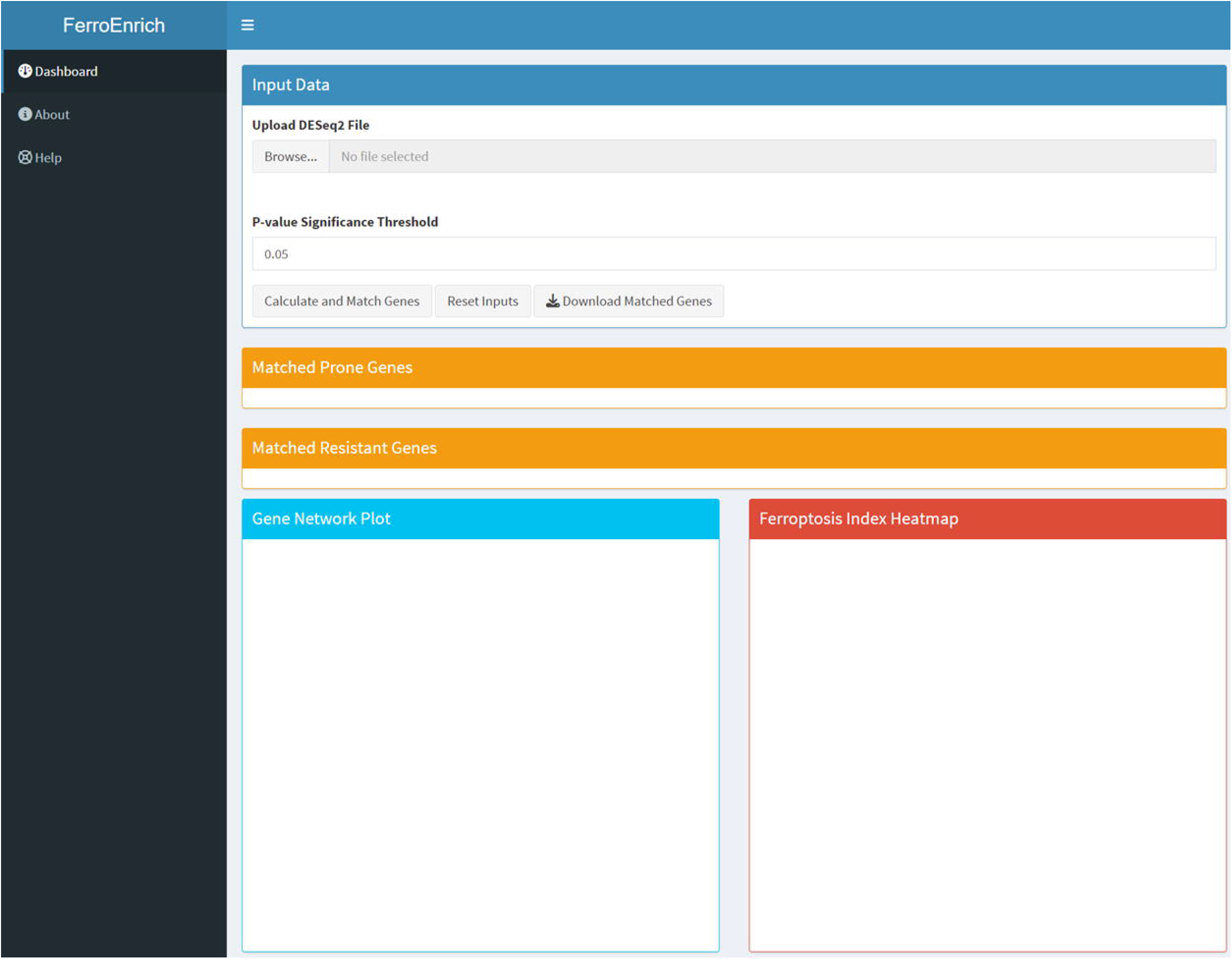
FerroEnrich Interface. This screenshot shows the user interface for the FerroEnrich web-based application. The interface allows users to upload DESeq2 gene expression data files and select p-value significant limits. Once the data has been submitted, users can compute and see the ferroptosis-prone and resistant genes. Additional capabilities include the option to download the matching genes and view them in a gene network diagram. The bottom panel shows the Ferroptosis Index Heatmap, which is a color-coded representation of ferroptosis potential based on gene expression data, allowing users to quickly assess and compare ferroptosis levels across several samples.

### 2.2. Data input and validation

The FerroEnrich procedure begins when users upload a DESeq2 file including gene expression data and associated metadata. The platform then validates the initial data to ensure that the file is complete and properly formatted. This validation is critical for avoiding problems in later analysis steps, as it checks for missing values, improper data types, and formatting discrepancies. Ensuring data integrity at this stage is critical for the accuracy, efficiency, and reliability of ferroptosis gene enrichment analysis results, allowing for a more robust downstream processing experience.

### 2.3. Gene identification and filtering

FerroEnrich analyses the data to identify the genes linked to ferroptosis after they have been verified. It makes use of predetermined standards derived from variations in gene expression:

- Prone genes: These are genes that have been found to be associated with the increased susceptibility of ferroptosis; they are distinguished in the dataset by positive Log2 fold changes accompanied by statistically significant p-values, indicating their overexpression.
- Resistant genes: On the other hand, negative Log2 fold changes, which denote downregulation, are indicative of genes that may impede ferroptosis.

### 2.4. Gene matching

Significant genes found in the gene expression data fetch with the database of prone and resistant genes and summarize the list of matched prone genes and resistant genes in the input data assigning ferro score for each gene to quantify its contribution to ferroptosis. Additionally, FerroEnrich improves user exploration by the creation of dynamic linkages to external databases such as STRING, a database devoted to network research and protein-protein interactions. This tool expands the scope of studies connected to ferroptosis by enabling researchers to dig further into the relationships, functions, and pathways associated with identified genes (Szklarczyk et al., 2019; 2021; 2023). This tool expands the scope of studies connected to ferroptosis by enabling researchers to dig further into the relationships, functions, and pathways associated with identified genes.

### 2.5. Gene Network Plot

The Gene Network Plot visualizes the relationships between the discovered ferroptosis-prone and resistant genes. Users can choose certain genes to examine how they interact with others in promoting or preventing ferroptosis. This plot aids in visualizing complicated interactions and can be an effective tool for hypothesizing regulatory pathways and prospective treatment targets.

### 2.6. Calculation of Ferroptosis Index value (FIV)

The primary analytical capability of FerroEnrich is its ability to measure a sample’s ferroptosis potential by calculating the Ferroptosis Index Value (FIV) by following these specific steps:

i. *Weighted Expression Calculation:* The FerroScore is a factor that represents the significance and contribution of each gene found in the analysis to ferroptosis and retrieved from FerrDb It is used to weight the expression of each gene. The following formula is used to determine each gene’s weighted expression (Wi) Calculation of Weighted Expression for each gene:

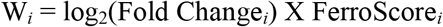

log2(Fold Changei): This denotes the log2-transformed fold change in gene i expression, offering an indication of the degree of change in gene expression in comparison to a control condition. FerroScorei Each gene has been given a score that represents its significance in the ferroptosis process, based on actual data or past knowledge.
ii. *Weighted Expression Summation*: For prone and resistant genes, separate summations are carried out. For prone genes (promoting ferroptosis):

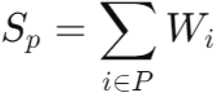

Here, Sp is the sum of weighted expressions for all prone genes, where P represents the set of genes that promote ferroptosis. This sum reflects the total promotive influence on ferroptosis in the sample. For resistant genes (resisting ferroptosis):

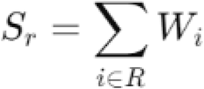

Similarly, Sr is the total of all resistant genes’ weighted expressions, where R is the set of genes that resist ferroptosis. The overall resistive influence against ferroptosis is represented by this sum.
iii. *Raw FIV Computation:* The total weighted expression of resistant genes is subtracted from that of prone genes to obtain the raw FIV. The Raw Ferroptosis Index Value is Calculated:

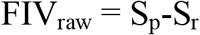
iv. *Scaling of FIV:* To standardize the index across various samples and situations, the raw FIV is scaled to a range of 0 to 5. Scaling of Raw FIV:

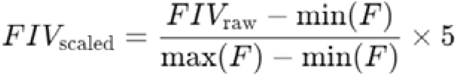
v. *Categorization:* The scaled FIV is divided into discrete levels (Normal, Mild, Moderate, High, Severe) for easier interpretation. This methodology ensures that the FIV accurately reflects the biological complexity of ferroptosis by including both promoting and resistive effects in a quantitative manner.

### 2.7. Ferroptosis Index Heatmap

The Ferroptosis Index Heatmap is an advanced, color-coded visual representation of the Ferroptosis Index Value (FIV). The hues on a spectrum ranging from green to red correlate to levels of ferroptosis potential: green for Normal, light green for Mild, yellow for Moderate, orange for High, and red for Severe as shown in **Figure 2**. This gradient scale provides a simple visual indication for quickly assessing ferroptosis risk across varied samples or situations. The colour and category of scaling as follows

**Figure 2:**
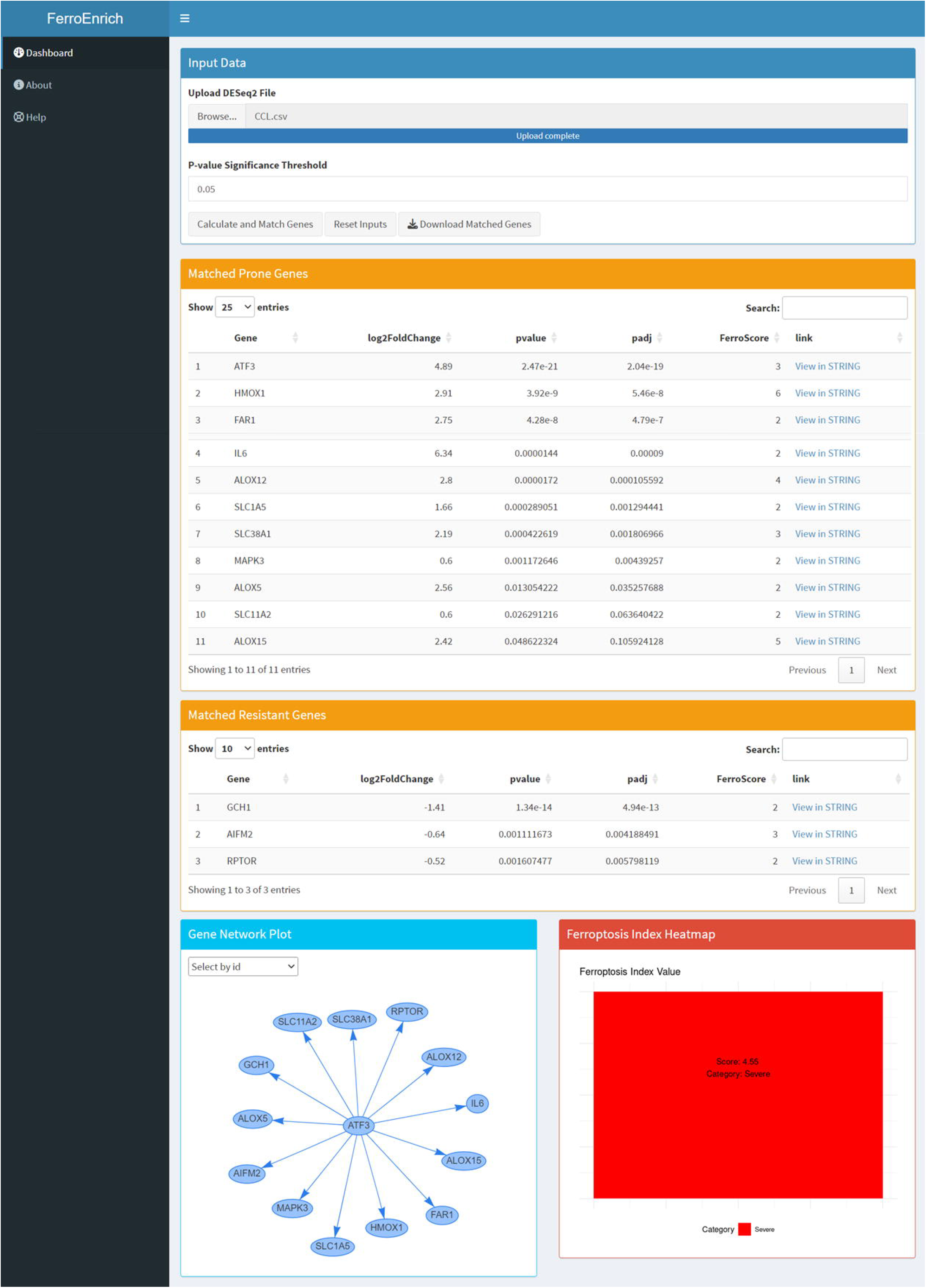
Output View of the FerroEnrich Tool. This graphic shows the results produced by the FerroEnrich tool after analyzing DESeq2 gene expression data. The interface parts featured are ‘Matched Prone Genes’ and ‘Matched Resistant Genes,’ which allow users to view a list of genes that promote or resist ferroptosis, respectively. In addition, the ‘Gene Network Plot’ visualizes the relationships between these genes, suggesting putative regulatory networks involved in ferroptosis. The ‘Ferroptosis Index Heatmap’ at the bottom provides a visual summary of ferroptosis potential across samples, classified by different levels of ferroptosis risk. This comprehensive output helps researchers understand intricate gene connections and the control of ferroptosis in biological samples.

Normal: FIV <= 2.50, shown in green, represents a normal ferroptosis potential. Mild: FIV values ranging from 2.50 to 2.99, indicated in light green, indicate a mild ferroptosis risk.

Moderate: FIV between 3.00 and 3.49, shown in yellow, indicates a moderate amount of ferroptosis.

High: FIV between 3.50 and 4.49, shown in orange, suggests a strong ferroptosis potential.

Severe: FIV of 4.50 or higher, shown in red, indicates significant ferroptosis potential.

## 3. Case Study and Examples

### 3.1. Study design

In order to evaluate the accuracy of our tool, we utilized the gene expression data of liver samples from humans at various fibrosis stages of non-alcoholic steatohepatitis (NASH) progression and data was retrieved from the GEO using the accession no GSE135251 (Govaere et al., 2020). In addition, we also used our gene expression data of Carbon tetrachloride (CCL4) -treated and control mouse liver samples. The main objective was to employ FerroEnrich, a web-based tool, to analyze and quantify the ferroptosis index (FIV) across these diverse experimental setups, thereby providing insights into the cellular mechanisms of liver fibrosis impacted by ferroptosis.

### 3.2. Results

The FerroEnrich method was used to analyze human liver samples at various fibrosis stages (F0-F4) of NASH (F0 to F4) of non-alcoholic steatohepatitis (NASH) and identified differential patterns in ferroptosis progression that corresponded to hepatic fibrosis severity. In the minimal fibrosis stage F0 and F1 ferroptosis-related gene expression changes were limited, with a Ferroptosis Index (FIV) of 2.73 and 2.89 indicating a mild risk of ferroptosis.

Fibrosis progression in stages F2 and F3 is marked by considerable ferroptosis, with FIVs of 3.02 and 3.23, respectively. These stages show increase of ferroptosis-prone genes such ELAVL1, TF, MAPK1, MAPK3, ACO1, TP53, BAP1, and STING1, as well as downregulation of resistance genes like GCLC, NFE2L2, GCH1, SLC7A11, and FXN. The F2 stage reveals early oxidative stress, but progressive fibrosis at F3 exhibits considerable changes, including marked overexpression of genes such as ELAVL1, MAPK3, and activation of stress markers TP53 and BAP1, resulting in a FIV of 3.23. The most advanced stage of cirrhosis (F4) is marked by strong expression of key ferroptosis-prone markers such as ACSL4, TF, MAPK3, STING1, and resistance genes GCLC, NFE2L2, GCH1, SLC7A11, and FXN, which corresponds to the highest FIV score of 3.68, indicating significant ferroptosis potential.

Significant variations in gene expression and FIV scores were detected in mice models treated with CCl4 to promote ferroptosis when compared to controls. FIV levels in treated animals increased significantly to 4.55, which was accompanied by the activation of stress response and lipid peroxidation genes such as ATF3, HMOX1, and ALOX12, as well as resistance markers SLC7A11, GCH1, and AIFM2. In contrast, control animals showed no consistent expression of ferroptosis-related genes and had a FIV of 0.0, indicating a normal liver state **(Table 1)**.

**Table 1.**
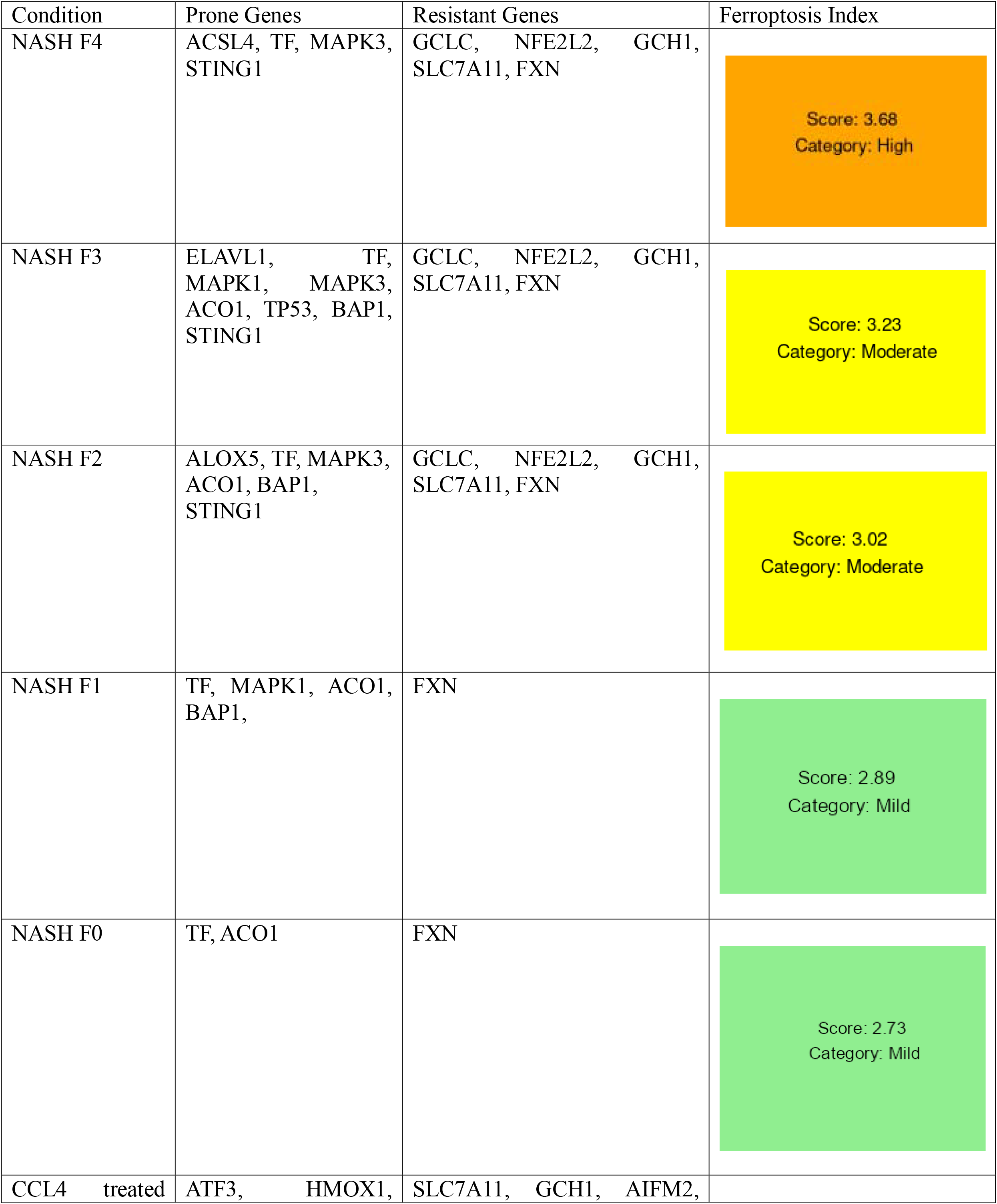
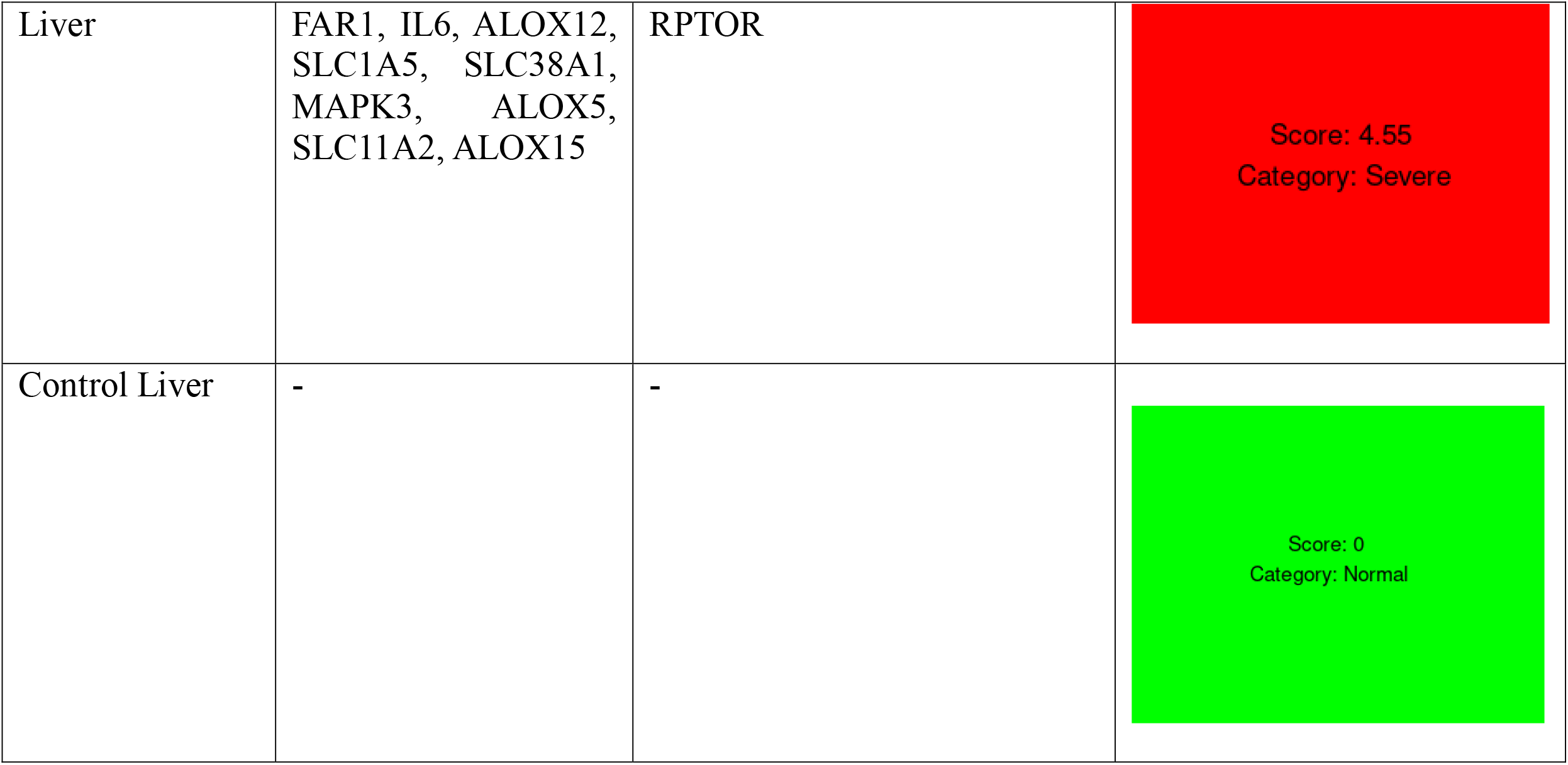
Compares ferroptosis-related gene expression and Ferroptosis index value at various phases of NASH progression in human liver samples and CCl4-treated mice models. Human results obtained utilizing the FerroEnrich method from samples at stages F0 to F4 of NASH (minimal to cirrhosis) reveal increasing increases in the Ferroptosis Index (FIV), with significant gene expression alterations indicating increased oxidative stress and lipid peroxidation in later stages. Concurrently, CCl4-treated mouse liver samples show significant overexpression of ferroptosis and stress response genes, which contrasts markedly with the control group. Data sources include GEO accession no. GSE135251.

These findings illustrate a clear correlation between the severity of liver fibrosis and increased ferroptosis potential, underscoring the role of ferroptosis in liver disease progression. The data from both human and mouse models not only highlight the consistency of ferroptosis mechanisms across different species but also reinforce the potential therapeutic implications of targeting ferroptosis in liver diseases, including NASH. This comprehensive case analysis provides a valuable foundation for further research into ferroptosis-based therapeutic strategies and underscores the importance of understanding the contributions of ferroptosis to the molecular dynamics of liver disease progression.

## 4. Discussion

The use of the FerroEnrich web application in this study demonstrates a substantial progress in the quantification and understanding of ferroptosis, particularly in the setting of liver disease such as NASH. FerroEnrich’s capacity to analyze and show extensive gene expression data on ferroptosis lays the groundwork for both basic research and possible clinical applications. The results provide strong evidence of the tool’s precision and reliability in discriminating between distinct phases of liver fibrosis and assessing the severity of ferroptosis at each stage.

FerroEnrich provides comprehensive outputs that enable a detailed study that associates particular gene expressions with ferroptotic reactions. Among these outputs are the Ferroptosis Index Values (FIV) for each sample, which is especially helpful in the setting of liver fibrosis, since alterations in cellular oxidation processes and iron metabolism are directly linked to the disease’s progression. FerroEnrich’s capacity to identify minute changes in gene expression associated with ferroptosis provides researchers with an extremely useful tool for precisely evaluating the severity and course of the disease.

The usage of FerroEnrich in this study demonstrates the tool’s strong analytical framework, which facilitates the integration of large datasets and provides practical insights into the molecular mechanisms behind ferroptosis. This accuracy not only improves our understanding of ferroptosis, but also helps to find possible biomarkers for early diagnosis and progression tracking of liver disorders. The results demonstrate FerroEnrich’s ability to handle and effectively evaluate high-throughput data, making it an invaluable tool in the study of cellular death pathways and their consequences in various illnesses.

Furthermore, FerroEnrich’s demonstration of consistent outcomes in mouse and human models lends additional credence to the tool’s efficacy and adaptability to various species and experimental configurations. This uniformity improves the validity of the conclusions drawn from such investigations by facilitating the translation of data from model organisms to human situations.

## 4. Conclusion

In conclusion, FerroEnrich has proven to be a highly accurate tool for assessing ferroptosis-related gene expressions, which makes it a great resource for studying the dynamics of gene expression in liver disorders and other conditions affected by ferroptosis. Researchers can gain precise insights into the cellular mechanisms underpinning disease progression thanks to FerroEnrich’s accuracy in mapping gene expression patterns and quantifying the ferroptosis index. This precision increases trust in the biological insights obtained from the data and strengthens the validity of applying these insights to guide clinical judgments and future directions in research. FerroEnrich’s creation represents a major advance in the field of ferroptosis research bioinformatics tools, providing a strong platform for further investigations into the intricate molecular landscape of ferroptosis. The potential influence of this tool on clinical diagnostics and academic research is substantial, and it could lead to the development of novel therapeutic strategies aimed at decreasing ferroptosis in chronic liver disorders and other conditions as it continues to be refined and adopted.

## Acknowledgments

This work was supported in part by start-up funds from the New York University Langone Hospital-Long Island, R01DK112971, R01DK135630, R01DK132056

## Competing interests

Authors declare that they have no competing interests.

## Notes

### Competing Interest Statement

The authors have declared no competing interest.

